# Proteomic analysis of isolated nerve terminals from Na_V_1.9 knockout mice reveals pathways relevant for neuropathic pain signalling

**DOI:** 10.1101/2024.06.28.601159

**Authors:** Ankita Rawat, Duc Tung Vu, Christoph Erbacher, Christian Stigloher, Nurcan Üçeyler, Matthias Mann, Michael Briese, Michael Sendtner

**Author notes:** Corresponding authors. Address: Institute of Clinical Neurobiology, University Hospital Wuerzburg, Versbacherstr. 5, 97078 Wuerzburg, Germany. E-Mail addresses (M. Briese); (M. Sendtner).

## Abstract

Neuropathic pain substantially affects the mental and physical well-being of patients and magnifies the socio-economic burden on the healthcare system. It is important to understand the molecular mechanisms underlying chronic pain to effectively target it. To investigate peripheral mechanisms relevant to pain signaling, we isolated nerve terminals from mouse footpads. The isolated peripheral terminals contain both pre- and post-synaptic proteins and are deficient in keratin and histone in both mice and humans. We detected the protein translational machinery and mitochondria in nerve terminals and observed that they were capable of endocytosis. An unbiased proteomic analysis of nerve terminals from footpads of Na_V_1.9 knockout mice shows dysregulation of the p38 mitogen-activated protein kinase (MAPK) and extracellular regulated kinase 1/2 (ERK1/2) pathways, and of protein components involved in translation and energy metabolism. Isolation of human nerve terminals from skin punch biopsies, validated by proteomic analysis, highlights the broad and translational value of our approach. Our study thus reveals peripheral signaling mechanisms implicated in pain perception.

## Introduction

Chronic pain, a debilitating condition affecting millions worldwide, comprises both persistent and recurrent pain sensations that last beyond the expected time for tissue healing, often leading to profound physical and psychological distress [31; 34]. The transition from acute to chronic pain involves a complex interplay of neurobiological processes in nociceptive sensory neurons located in dorsal root ganglia (DRG), including hyperexcitability of nociceptors, alterations in neuropeptide production and release, dysregulation of inflammatory mediators, and aberrant associations of nociceptive nerve terminals with the microenvironment in affected areas [30; 44]. While pain-signaling mechanisms have been shown to alter gene expression and protein production in somas of DRG sensory neurons, much less is known about proteome alterations occurring in peripheral nerve terminals in the skin in pain contexts.

The peripheral branches of DRG sensory neurons innervates the epidermis and dermis [45] where they are in contact with multiple cell types. Nociceptive signaling is influenced locally by such non-neuronal cells like macrophages, immune cells, nociceptive Schwann cells and keratinocytes, which play an important role in modulating pain thresholds and sensitivity within the emerging concept of cutaneous nociception [24; 42]. Keratinocytes are the predominant cells in the epidermis and are involved in direct communication with the peripheral sensory nerve forming the neuro-cutaneous unit [16; 51]. They are also involved in signal transduction and nociception [2; 48; 50].

In this study, we isolated peripheral nerve terminals from the mouse footpad using a protocol that was previously applied for synaptosome isolation from the central nervous system [4]. We observed the presence of several pre- and postsynaptic proteins in isolated nerve terminals and performed mass spectrometry to identify the proteome of these terminals. Further, we analysed nerve terminals of Na_V_1.9 knockout mice, which show impaired mechanical and thermal sensory capacities along with diminished electrical excitability of nociceptors [19]. This identified translation and mitochondrial energy metabolism as well as signaling cascades such as the p38 mitogen-activated protein kinase (MAPK) and extracellular regulated kinase 1/2 (ERK1/2) appears particularly relevant for pain perception. Additionally, the isolation of human nerve terminals from skin punch biopsies, as validated by proteomic analysis, underscores the extensive and translational significance of our approach. Therefore, our data reveals the proteomic diversity of peripheral nerve terminals, and the techniques outlined here could be applied to study local protein alterations in models of chronic pain, pain resolution as well as in diagnostic human biomaterial.

## Results

### Isolation of enriched peripheral nerve terminals from mouse footpads

As a starting point, we investigated the presence of synapse-like contacts in the epidermis of the mouse footpad. To assess this, we performed vibratome sectioning of footpad tissue and immunostained sections for neurofilament heavy chain (Nefh) and the synaptic markers postsynaptic density protein 95 (Psd95) and synapsin 1 (Syn1). We observed co-localisation of the synaptic markers with Nefh, which was further confirmed with high-resolution Structured Illumination Microscopy (SIM) (Fig. 1A and B). This observation thus indicates the presence of synaptic components in the epidermis of the mouse footpad.

**Figure 1.**
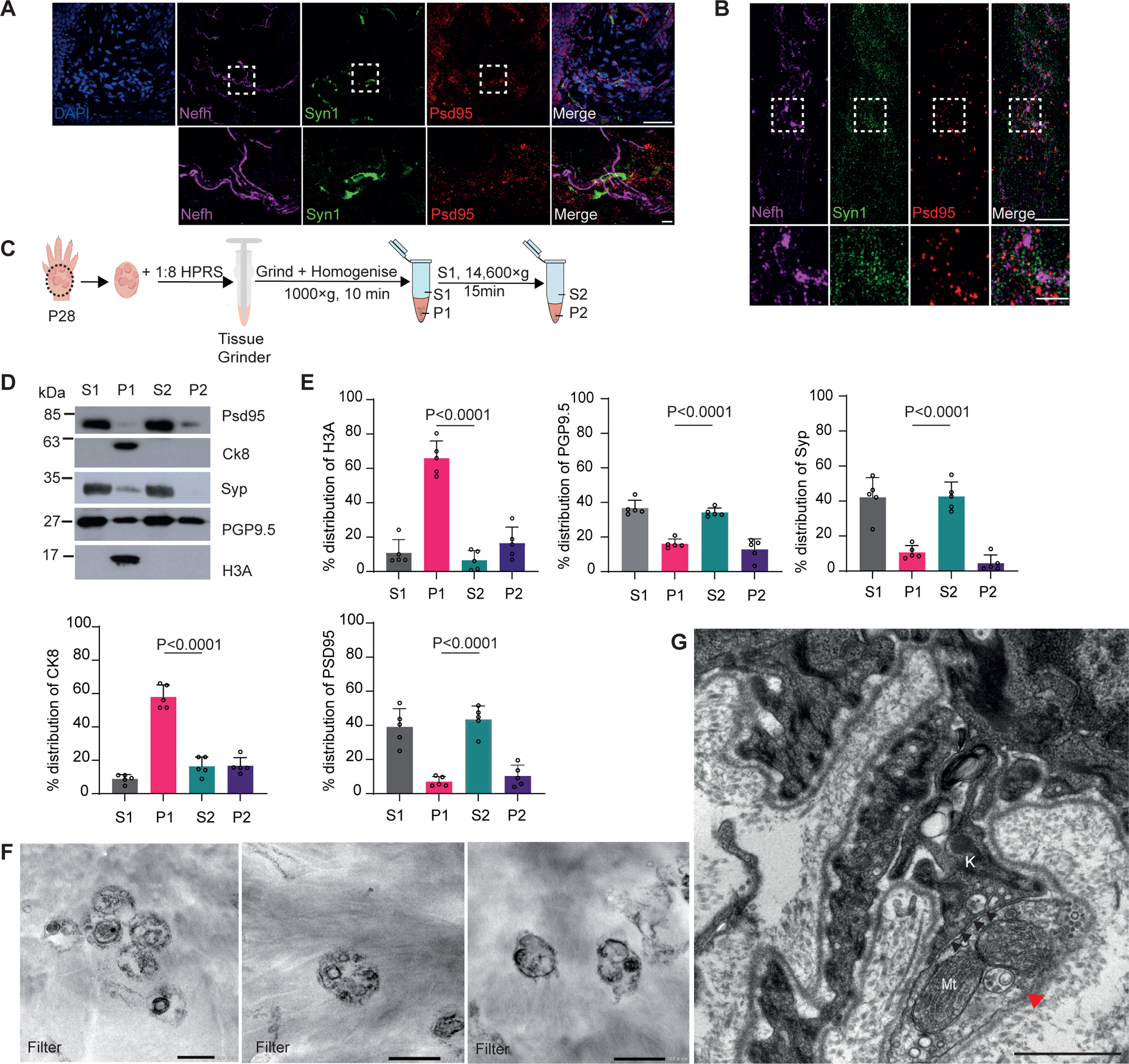
Isolation of peripheral terminals from mouse footpad. (A and B) Confocal and SIM images of mouse epidermis stained with Nefh, Syn1 (Synapsin 1), Psd95 (post-synaptic density 95), and DAPI show a synapse-like structure. Scale bars: 50 µm and 5 µm (magnified areas). (C) Schematic showing peripheral nerve isolation from P28 mouse footpad by subcellular fractionation using HPRS: HEPES protease RNAse, and sucrose buffer. (D) Representative immunoblot of enrichment of synaptic markers, PSD95, Syp1, and PGP9.5 in the S2 fraction. (E) Quantification of synaptic and nuclear proteins in different fractions after subcellular fractionation in (D) One-way Anova with Tukey’s multiple comparisons tests. Data are meanL±Ls.d. of nL=L5 biological replicates. (F) Representative image of mouse footpad epidermis showing contact site (black arrowheads) between peripheral nerve terminal (red arrowhead), and keratinocyte (K), Mitochondria (Mt). Scale bar: 500 nm. (G) Representative electron micrographs of isolated peripheral terminals following fractionation immobilised on polycarbonate filter. Scale bar: 200 nm.

Next, we sought to biochemically isolate nerve terminals from mouse footpads. Methods for synaptosome isolation from the brain were first introduced by Whittaker et al. [57]. In our current study, we have adapted this technique to isolate peripheral nerve terminals from the mouse footpad. This area, densely populated with pain-sensing nerve terminals [6], has not been previously explored using this isolation technique. We lysed footpad tissue and performed subcellular fractionation by differential centrifugation to obtain supernatant (S) and pellet fractions (P) (Fig. 1C). We observed that the S2 fraction contained the synaptic markers Psd95 and synaptophysin (Syp) as well as the axonal marker protein gene product-9.5 (PGP9.5) and lacked the nuclear marker histone H3A and the keratinocyte marker cytokeratin 8 (Ck8) (Fig. 1D and E). This indicates that the S2 fraction contains nerve terminals. We next examined the ultrastructure of the peripheral nerve terminals in the epidermis by Transmission Electron Microscopy (TEM). This revealed a terminal adjacent to the basal membrane and in contact with a process of a basal epidermal cell that could be a keratinocyte (Fig. 1G). The contact zone between the terminal and the possible keratinocyte process seemed to show specializations. The contacting membranes appeared straight, and the cleft contained fibrous material resembling that found in synaptic clefts in the CNS, possibly indicating contact proteins and vesicles (Fig. 1G, Supplementary Fig. 1A). We then examined the structure of isolated peripheral nerve terminals by TEM following immobilization of the isolated peripheral nerve terminals on polycarbonate membrane filters. Interestingly, we observed structures similar to synaptosomes isolated from CNS-like brain (Fig. 1F, Supplementary Fig. 1B) [4; 15]. This led us to further investigate the S2 fraction containing the peripheral nerve terminals of sensory neurons.

Next, we sought to confirm that nerve terminals of nociceptive neurons can be enriched with this isolation method. For this purpose, we crossed Na_V_1.8cre mice [1] with CAG-CAT-eGFP [35] mice to activate GFP expression selectively in nociceptors. Na_V_1.8 is a key voltage-gated sodium channel present predominately in the peripheral nociceptive neurons [3] and plays a significant role in pain perception [62]. Na_V_1.8 expression is also upregulated in patients with chronic neuropathic pain [12; 53] and animal models of inflammatory pain [56]. Na_V_1.8 blockers and knockdown studies lead to an attenuation of pain associated behavior in neuropathic pain models [23; 61]. Immunohistochemistry of the footpad epidermis of control Na_V_1.8cre and Na_V_1.8cre::CAGeGFP mice revealed GFP-positive fibers in the Na_V_1.8cre::CAGeGFP footpads which were absent in the control (Fig. 2A). The eGFP-positive fibers co-localized with the axon marker PGP9.5, which reflects the dense innervation of mouse epidermis by nociceptive fibers (Supplementary Fig. 2). We then isolated peripheral nerve terminals as before and observed that GFP was present in the S2 fraction together with the synaptic markers (Fig 2B and C). This indicates that the S2 fraction contains isolated peripheral nerve terminals from nociceptors that could be further characterized in this study.

**Figure 2.**
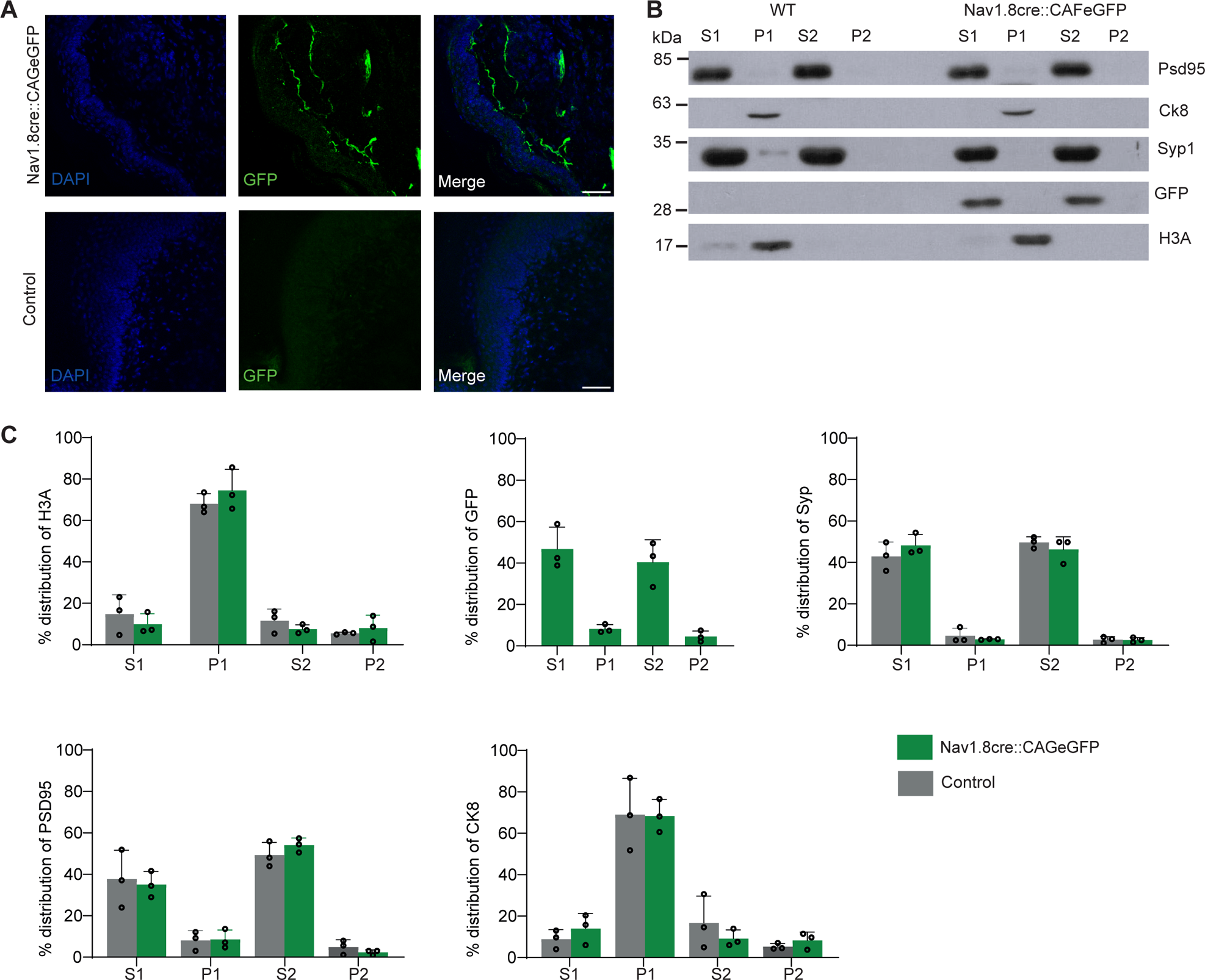
Presence of nociceptive nerve terminals during fractionation. (A) Immunohistochemical detection of Na_V_1.8 positive fibers using an antibody against GFP and DAPI in the longitudinal epidermal section of P28 wild type and Na_V_1.8cre::CAGeGFP mice. Scale bar 50 µm. (B) Representative immunoblot of enrichment of synaptic markers, PSD95, Syp, and GFP in the S2 fraction. (C) Quantification of synaptic, nuclear, and GFP protein in different fractions after subcellular fractionation in (B). Data are meanL±Ls.d. of nL=L3 biological replicates.

### Isolated peripheral nerve terminals contain active protein translational machinery

To characterize the isolated peripheral nerve terminals, we immobilised them on coverslips following a method previously introduced by Ramirez et al. [55] with minor adjustments. With this approach, we observed that approximately 65% of the particles contained both pre- and post-synaptic structures as they co-stained for Nefh, Psd95, and Syn1 (Fig 3A and B). Previous studies have shown that peripheral nerve endings contain mitochondria and ribosomes [26; 47]. To test whether this is also the case for our isolated nerve terminals, we stained them for ribosomal markers Rps6 and Rpl24, and the mitochondrial markers Tomm20, respectively, along with Syp. The percentage co-localisation between the ribosomal and synaptic markers was 87.5% (Fig 3C and D) and between the mitochondrial and synaptic markers 75.8% (Fig 3E and F). This finding indicates the presence of the protein translation machinery and mitochondria for local energy supply in the isolated peripheral nerve terminals. To evaluate the functionality of peripheral nerve terminals, we incubated the S2 fraction with the membrane dye mCLING, which can be used to monitor membrane trafficking [41]. Stimulation of peripheral nerve terminals with KCl induced endocytosis of mCLING (Fig. 3G). As a control for specificity, endocytosis of mCLING was not observed when the temperature was shifted to 4°C, a condition that is known to abolish active cellular endocytosis (Fig. 3G). This indicates that the isolation procedure preserves the functionality of peripheral terminals.

**Figure 3.**
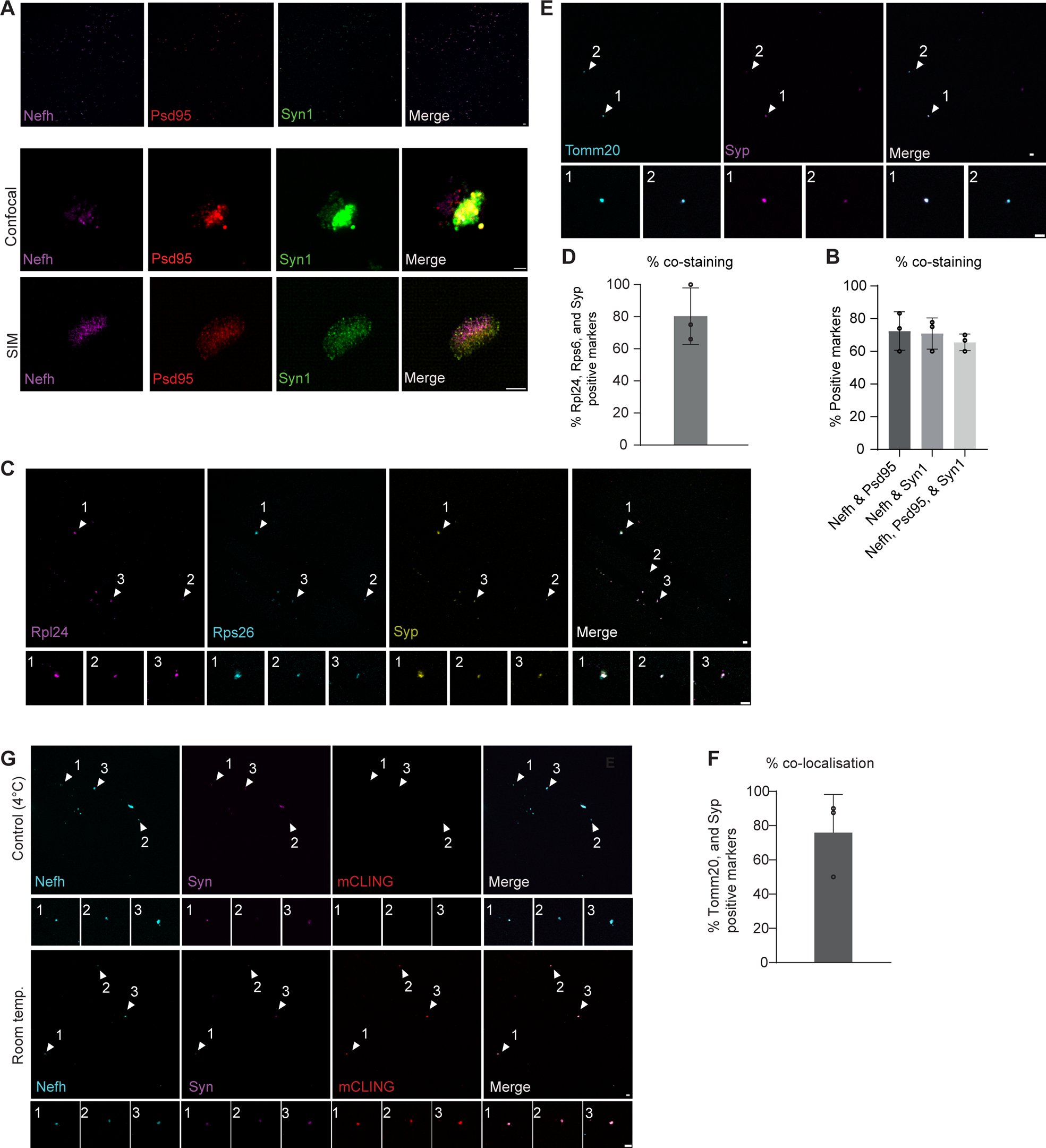
Isolated peripheral terminals contain mitochondrial and ribosomal components. (A) Representative images of immobilised peripheral nerves on coverslips and stained with Nefh, Psd95, and Syn1 (confocal 60x, confocal digital zoom, and SIM respectively). (B) Quantification of percent co-staining of Nefh and Psd95, Nefh and Syn1, and Nefh, Psd95, and Syn1 of (A). Data are meanL±Ls.d. of nL=L3 biological replicates. (C) Representative images of immobilised peripheral nerves on coverslips and stained with ribosomal markers RPL24 and S6 along with Synaptophysin. Data are meanL±Ls.d. of nL=L3 biological replicates. (D) Quantification of percent co-staining of ribosomal and synaptic markers of (C). Data are meanL±Ls.d. of nL=L3 biological replicates. (E) Representative images of immobilised peripheral nerves on coverslips and stained with mitochondrial marker, Tomm20 and Syp. Data are meanL±Ls.d. of nL=L3 biological replicates. (F) Quantification of percent co-staining of Tomm20 and Syp in (E). Data are meanL±Ls.d. of nL=L3 biological replicates. (G) Representative images showing the mCLING dye being endocytosed by the peripheral nerve endings at room temperature and absence of mCLING dye in the peripheral nerve endings at control (4°C). Scale bar: 5 µm.

### Isolated peripheral terminals are devoid of keratins and chromatin in mouse skin

To further assess the purity of the isolated peripheral terminal from previous experiments, we conducted Mass Spectrometry (MS)-based proteomics of the S2 fraction. As a control, we utilized the P1 fraction, which contains markers for nuclei and cytoplasmic components of keratinocytes, previously indicated by the western blotting (Fig. 1D). As an initial step, we subjected the samples to SDS-PAGE followed by silver staining and observed similar patterns of bands between individual replicates for the P1 and S2 fractions indicating reproducibility of the isolation procedure (Fig. 4A). In agreement, principal component and heat map analysis showed clustering of replicates for each fraction (Fig. 4B and C).

**Figure 4.**
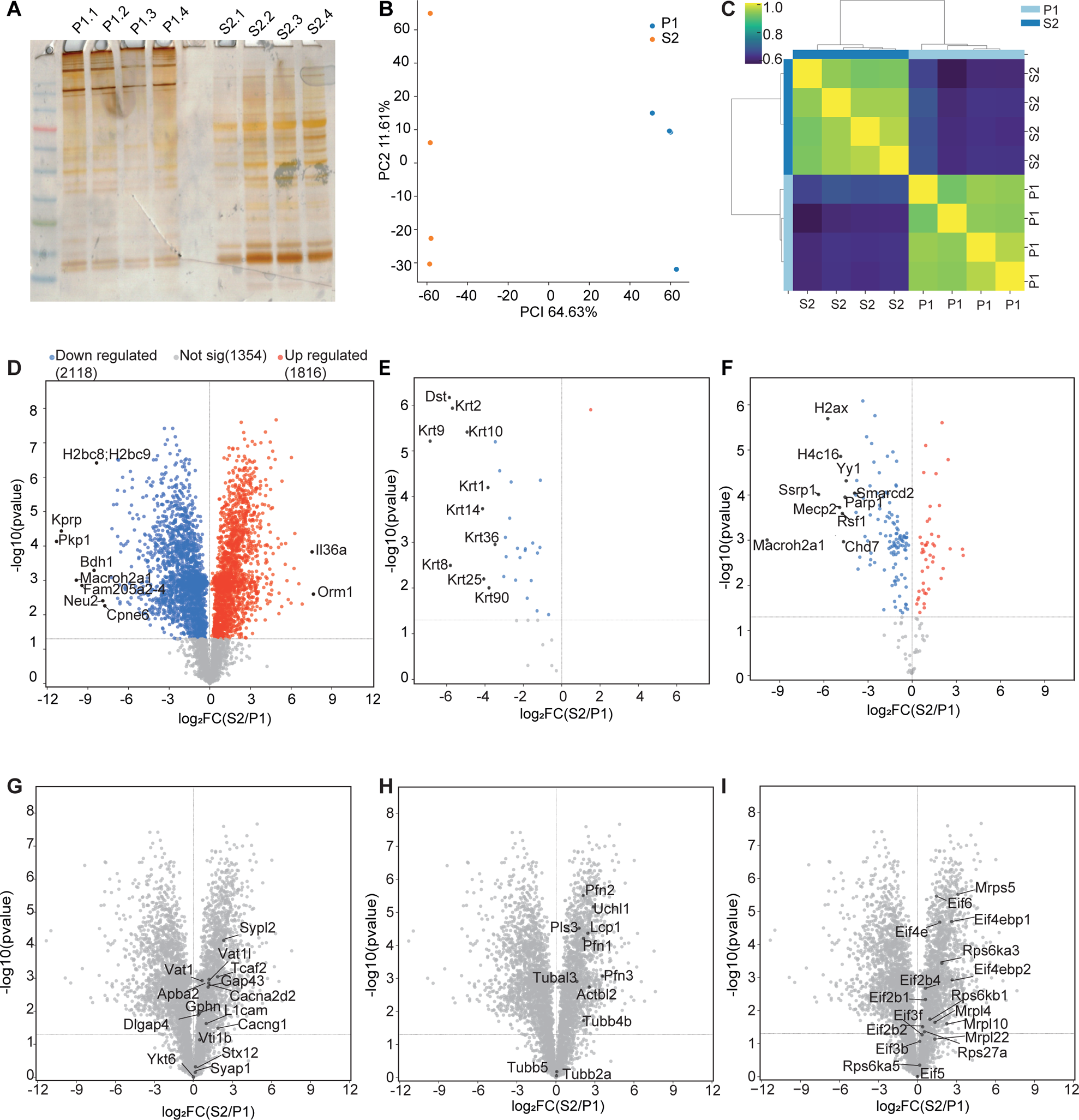
Isolated peripheral terminals from mouse are deficient in histone and keratin. (A) Silver staining of protein isolated from P1 fraction (lanes 1-4) and S2 fraction (lanes 5-8). (B and C) Principal component analysis and heat map with the Pearson correlation coefficient between samples showing high reproducibility of S2 and P1 fractions from n=4 biological replicates. (D) Volcano plot showing relative protein enrichment of S2 compared to P1 fractions. The red and blue dots depict significantly (p<0.05) enriched proteins in S2 and P1, respectively. (E and F) Volcano plots of pre-defined keratin (GO term ‘keratin filaments’) (E) and chromatin-associated proteins (GO term ‘chromatin’) (F). (G-I) Volcano plot of the S2 fraction vs the P1 fraction with manual annotation for pre- and post-synaptic markers and ion channel components (G), cytoskeletal proteins (H) and ribosomal proteins and translation factors.

Protein enrichment analysis identified 1816 proteins that were enriched in the S2 fraction and 2118 proteins that were enriched in the P1 fraction (p<0.05) (Fig. 4D). Importantly, proteins enriched in the P1 fraction contain keratin filaments and chromatin components (Fig. 4E and F), which is in line with the immunoblot analysis (Fig. 1D) and further validates the efficiency of the fractionation procedure. Importantly, many pre-synaptic proteins such as synaptophysin-like protein 1 (Sypl1), synaptophysin-like protein 2 (Sypl2), synapse-associated protein 1 (Syap1), and synaptic vesicle membrane protein VAT-1 homolog (Vat1) were present in the S2 fraction (Fig. 1G). Post-synaptic markers such as gephyrin (Gphn), disks large-associated protein 4 (Dlgap4), and neuromodulin (Gap43) were also detectable in the S2 fraction (Fig. 1G), which indicates the presence of both pre- and post-synaptic structures. While ion channels and other transmembrane proteins are usually less detectable by mass spectrometry, we observed voltage-dependent calcium channel subunit alpha-2/delta-2 (Cacna2d2), voltage-dependent calcium channel gamma-1 subunit (Cacng1), and TRPM8 channel-associated factor 2 (Tcaf2) in the S2 fraction (Fig. 4G). Surprisingly, regulators of actin dynamics such as profilin1, 2, 3 and plastin3 were also found in S2, which are important for synaptogenesis (Fig. 4H) [18; 33]. Finally, we observed components of ribosomes and translation factors in the S2 fraction (Fig. 4I), in agreement with previous observations that ribosome assembly is dynamically regulated in axon terminals [10]. This demonstrates that the isolated peripheral nerve terminals lack non-neuronal components such as keratinocytes and nuclei and are enriched in pre- and post-synaptic markers along with protein synthesis machinery in the S2 fraction, highlighting its suitability for subsequent experiments.

### Analysis of Na_V_1.9 knockout nerve terminals reveals pain signaling pathways

The voltage-gated sodium channel Na_V_1.9 is strongly expressed in nociceptors and plays a significant role in neuropathic and inflammatory pain [11; 29; 36]. Na_V_1.9 knockout (KO) mice, in which exons 4 and 5 of the *Scn11a* gene encoding Na_V_1.9 were replaced by a neomycin resistance cassette, have no Na_V_1.9-mediated sodium currents [37] and display compromised mechanical and thermal sensory functions, alongside diminished electrical excitability of nociceptors [19]. Moreover, patients with mutations in the gene for Na_V_1.9 display impaired pain perception [22; 27]. In order to detect proteins that are less expressed and thus correlate with reduced pain sensation, we performed proteomics analysis on S2 fractions from enriched peripheral nerve terminals from Na_V_1.9 KO and wild type mice (Supplementary 3A-D). Our analysis revealed 738 proteins enriched in wild type nerve terminals and 759 proteins enriched in Na_V_1.9 KO terminals (p<0.05) (Fig. 5A). One of the most strongly enriched proteins in wild type relative to Na_V_1.9 KO nerve terminals is protein phosphatase 1 E (Ppm1E), which is increased in inflammatory pain by regulation of MEK/ERK signaling in spinal cord dorsal horn of rats [20] (Fig. 5B). Interestingly, another one of the most strongly enriched proteins in wild type relative to Na_V_1.9 KO nerve terminals is Nefl (Fig. 5B). Nefl is upregulated upon spinal nerve ligation-induced neuropathic pain and repression of Nefl attenuates pain [60]. Functional classification revealed enrichment of proteins related to translation, monoatomic ion channels, synapse, sensory perception to pain, and response to pain (Fig. 5C).

**Figure 5.**
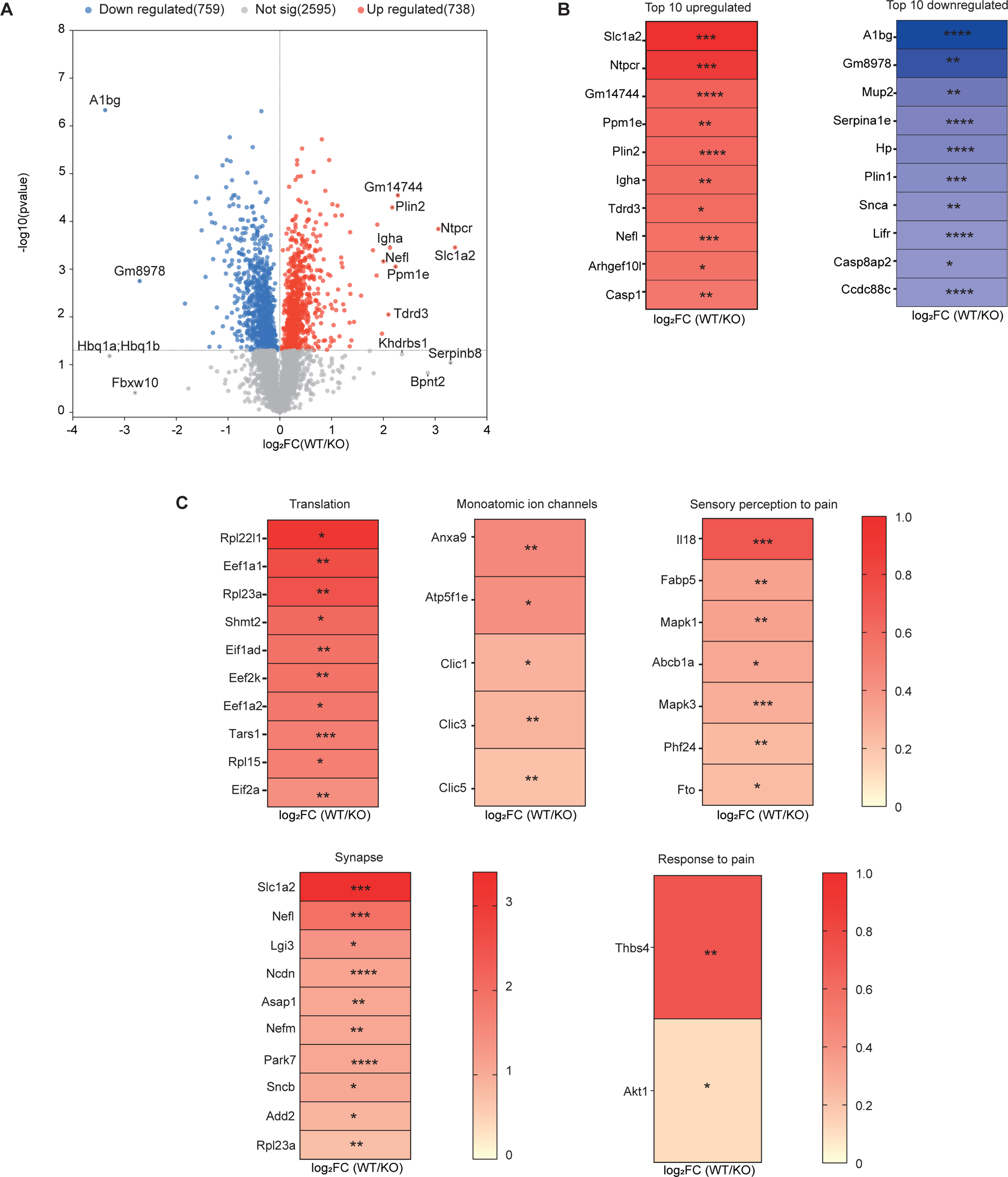
Protein enrichment analysis of wild type relative to Na_V_1.9 KO nerve terminals. (A) Volcano plot showing relative protein enrichment of the S2 fractions from wildtype compared to Na_V_1.9 KO mice. The red and blue dots depict the significantly (p<0.05) upregulated and downregulated proteins, n=3 biological replicates. (B) Heat maps showing the top 10 upregulated and top 10 downregulated proteins. (C) Differentially enriched proteins associated with the gene ontology categories translation, monoatomic ion channels, sensory perception to pain, synapse, ion channels, and response to pain. The annotation in the volcano plots is log2 fold change of top DEPs

Next, to associate differentially expressed proteins with gene function in an unbiased manner, we examined enrichment for Gene Ontology (GO) terms related to biological process, molecular functions, and cellular components, using DAVID (Fig. 6A and B; Supplementary Fig. 4A-D) [21; 43]. GO term enrichment was visualized with the SRplot tool [52], with a significance threshold set to p<0.05. Among the top 20 GO terms based on log2FC, proteins enriched in wild type relative to Na_V_1.9 KO nerve terminals, we observed ‘intermediate filament bundle assembly’, ‘Arp2/3 complex-mediated actin nucleation’, and ‘regulation of actin filament polymerization’ (Fig. 6A). Among the top 20 GO terms for proteins reduced in wild type relative to Na_V_1.9 KO nerve terminals, we observed GO terms related to energy metabolism and calcium ion transport (Fig. 6B).

**Figure 6.**
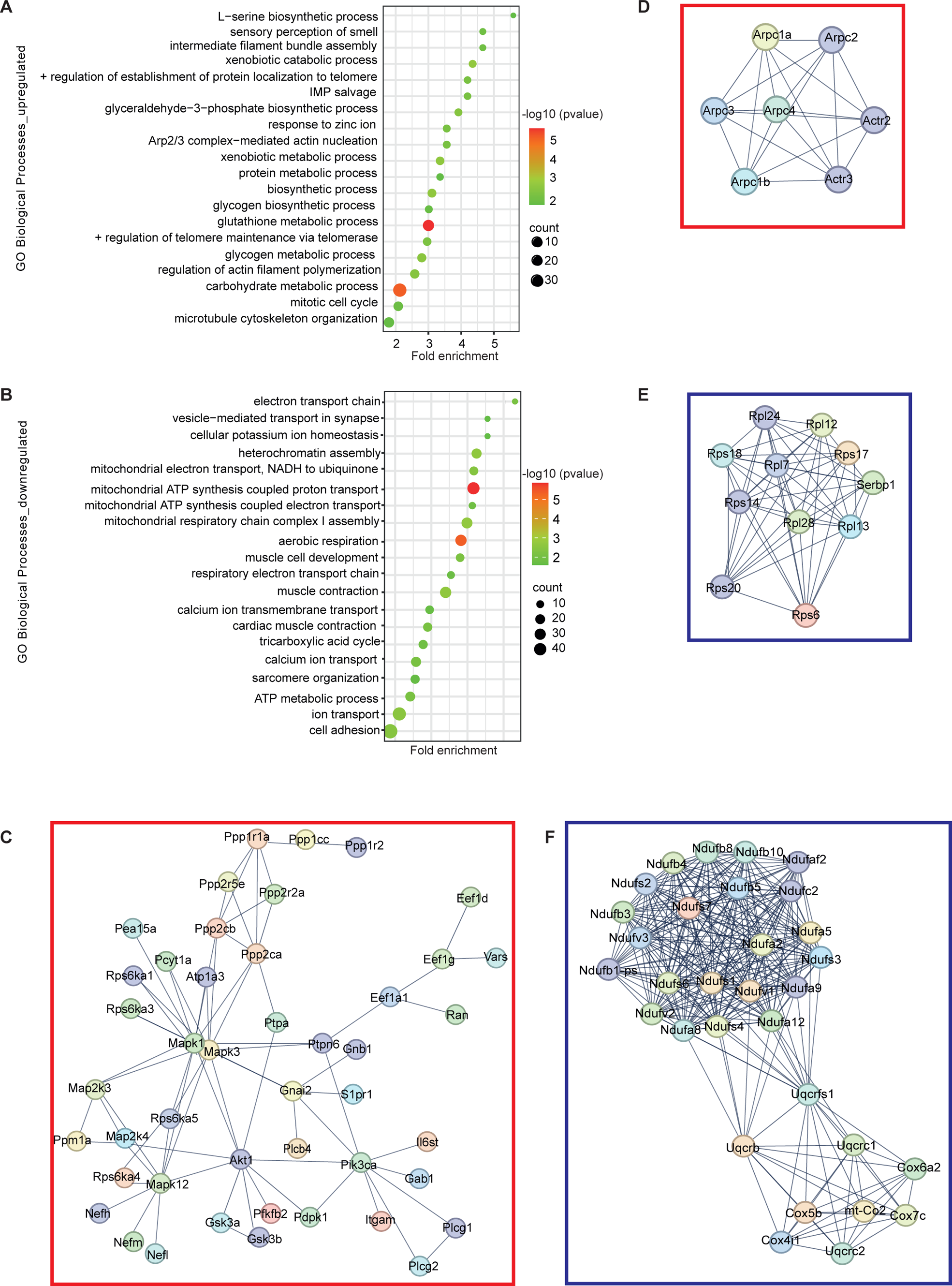
Identification of pain signaling pathways in nerve terminals. (A and B) Dot plot summarizing the top 20 Gene Ontology (GO) biological processes terms for upregulated (A) and downregulated (B) proteins in peripheral nerve terminals of wild type relative to Na_V_1.9 KO mice. The x-axis represents the fold enrichment, and the y-axis represents the enriched pathway for biological processes. The colour scale indicates different thresholds of the p-value, and the size of the dot indicates the number of genes corresponding to each pathway. (C-F) STRING analysis of proteins up- (C and D) and downregulated (D and E) in wild type relative to Na_V_1.9 KO nerve terminals. Each node represents a protein, and each edge represents an interaction including either physical or functional associations. Only interactions with the highest confidence score (0.9) are shown.

STRING analysis [49] was then employed to construct protein-protein interaction networks of the differentially expressed proteins. Importantly, components of the ERK1/2 and p38 MAPK pathways (Mapk1, Mapk3, Mapk12, Map2k3, Map2k4, and Mapk12) were increased in wild type relative to Na_V_1.9 KO nerve terminals (Fig. 6C, Supplementary 5). Since ERK1/2 and p38 MAPK are activated during peripheral inflammation in an NGF-dependent manner [8; 25], our result shows the significance of these pathways for pain signaling. Additionally, cytoskeletal proteins and regulators such as components of the Arp2/3 complex were also enriched in wild type nerve terminals (Fig. 6D, Supplementary 5), in agreement with the GO term analysis (Fig. 6A). In contrast, components of the translation machinery and mitochondria were reduced in wild type nerve endings, which agrees with observations that mitochondrial dysregulation can contribute to chronic pain [58] (Fig. 6E and F, Supplementary 6). In conclusion, our data show that MAPK pathways and cytoskeletal components are associated with nociception, whereas dysregulation of translation and energy metabolism might interfere with pain perception.

### Isolation of peripheral nerve endings from human skin biopsies

Finally, we applied the peripheral nerve isolation technique to healthy human skin biopsy-derived epidermis, separated from the dermal and hypodermal layers and conducted MS-based proteomics of the P1 and S2 fractions. Silver staining of size-fractionated protein samples showed similar band patterns between replicates, indicating reproducibility of the isolation procedure (Fig. 7A). This was corroborated by principal component and heat map analysis, demonstrating the clustering of replicates for each fraction (Fig. 7B and C). Protein enrichment analysis identified 860 proteins that were enriched in the S2 fraction and 4270 proteins that were enriched in the P1 fraction (p<0.05) (Fig. 7D). Similar to mouse skin samples, proteins enriched in the P1 fraction of human skin biopsies contained keratin filaments and chromatin components (Fig. 7E and F). Comparison of proteins enriched in P1 and S2 fractions showed a strong overlap between fractions from human skin biopsies with those obtained from mouse footpads (Fig. 7G). This demonstrates that peripheral nerve terminals can also be isolated from diagnostic human skin biopsies.

**Figure 7.**
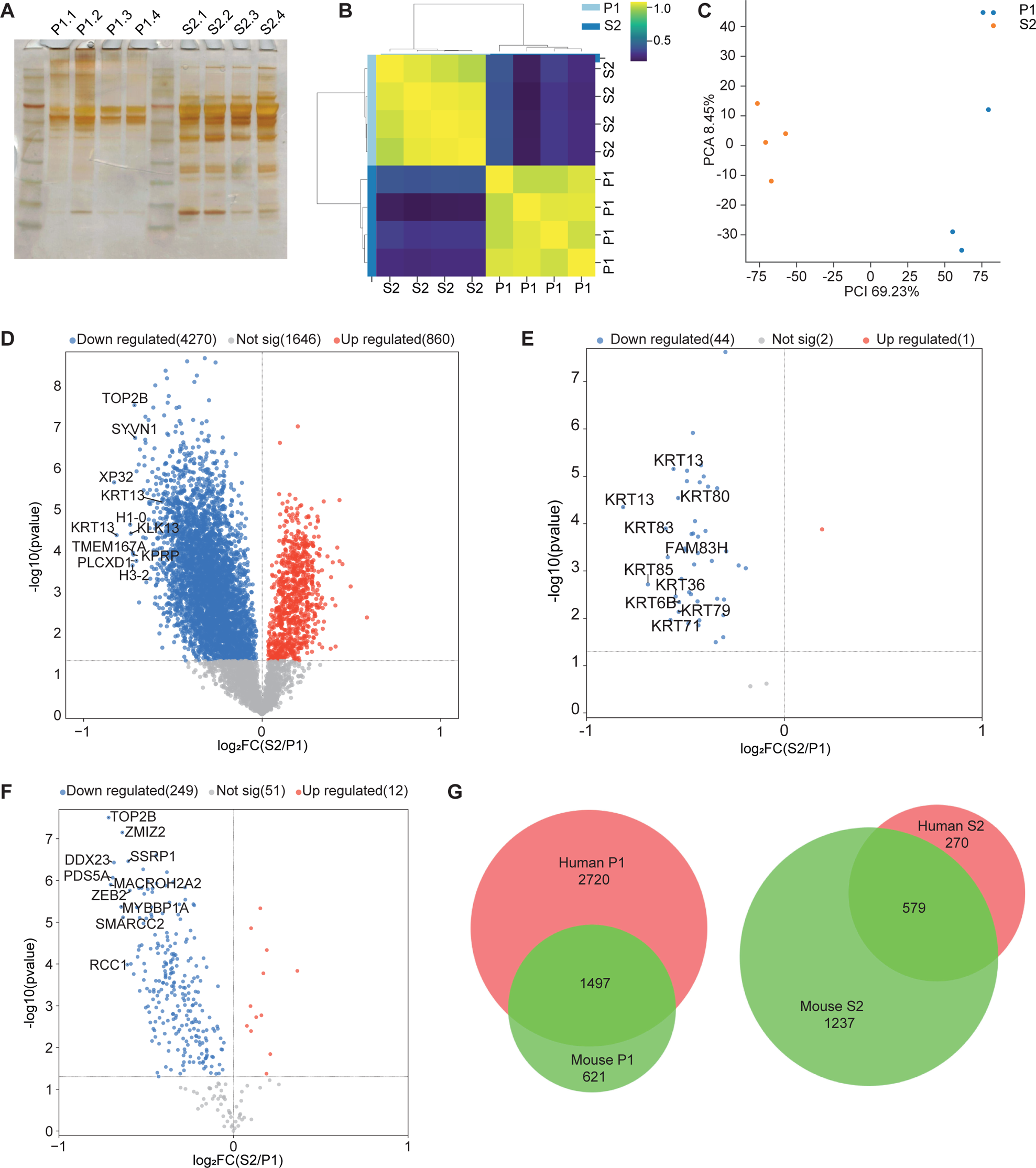
Isolated peripheral terminals from humans lack histone and keratin. (A) Silver staining of protein isolated from P1 fraction (lanes 1-4) and S2 fraction (lanes 5-8). (B and C) Principal component analysis (B) and heat map (C) with the Pearson correlation coefficient between samples showing high reproducibility of S2 and P1 fractions from n=4 biological replicates. (D) Volcano plot showing relative protein enrichment of S2 compared to P1 fractions. The red and blue dots depict significantly (p<0.05) enriched proteins in S2 and P1, respectively. (E and F) Volcano plots of pre-defined keratin (GO term ‘keratin filaments’) (E) and chromatin-associated proteins (GO term ‘chromatin’) (F). (G) Venn diagram representing the overlap in differentially enriched proteins in the P1 and S2 fraction of mouse and human samples.

## Discussion

Chronic pain is an incapacitating disorder representing a substantial health burden, affecting one in five people in Europe, and for half of these individuals, treatment is inadequate [5; 14]. While previous studies describe mechanisms underlying gene expression and protein translation alterations in sensory neurons during chronic pain [38], considerably less is understood about local proteome alterations in peripheral nerve endings at the neuro-cutaneous unit. The approach proposed here employs the isolation of peripheral nerve terminals from mouse footpad that are densely innervated with nociceptor terminals.

The immunohistochemistry results illustrated an overlap between the pre- and post-synaptic marker with Nefh, which was additionally confirmed by the presence of synaptic markers and PGP9.5 in the S2 fraction after fractionation. Interestingly, the electron micrograph of the whole epidermis suggests the presence of a close contact site between the peripheral nerve and keratinocytes, resembling an en passant synapse [51]. The observation of GFP in the S2 fraction from Na_V_1.8cre::CAGeGFP footpads provides evidence for the enrichment nociceptor terminals with the isolation method described here.

One of the most relevant findings of our study is the detection of both pre- and post-synaptic structures in the S2 fraction. This result indicates the presence of synapse-like contacts between sensory nerve terminals and epidermal cells, most likely keratinocytes. Signaling from keratinocytes to nociceptors has been shown before [2], indicating a modulatory function of skin cells for pain perception. More recently, contacts between keratinocytes and sensory afferents have also been observed at the ultrastructural level [16]. In line with previous studies [28; 32], we also detected components of the translational machinery and mitochondrial proteins in nerve terminals, underscoring their capacity for energy metabolism [13] and protein synthesis. This agrees with observations that ribosomes are not only present in the cell body but also in the terminals of sensory neurons [59]. Most importantly, the S2 fraction is also depleted of histone and keratin proteins for both mice and human skin samples, which further indicates the efficiency of the enrichment procedure.

Na_V_1.9 plays an important role in the induction of thermal and mechanical hypersensitivity, both in subacute and chronic inflammatory pain models. Hence, we applied an unbiased proteomics approach to identify proteins enriched in wild type relative to Na_V_1.9 KO terminals, as these proteins indicate pathways required for pain perception. Interestingly, we observed components of the ERK1/2 and p38 MAPK signaling pathways upregulated in wild type terminals. These pathways are activated during pain in an NGF-dependent fashion. NGF has been demonstrated to increase the expression of the heat-sensitive transient receptor potential (TRP) channel, TRPV1, via activation of the tyrosine kinase receptor (Trk) A/p38 MAPK pathway [25]. Additionally, we observed increased levels of Nefl in wild type relative to Na_V_1.9 KO nerve terminals. Elevated levels of Nefl have been detected in peripheral serum in patients with bortezomib-induced polyneuropathy (BIPN) [7], and these elevated levels might not only reflect enhanced release from degenerating neurons but also enhanced production in sensory neurons with enhanced pain perception.

We successfully adopted our peripheral nerve isolation technique to isolate and analyze human nerve terminal proteins from diagnostic skin biopsy samples. In the context of hardly available human sensory neuron and nerve biomaterial for research and diagnostics, we present an innovative, minimally invasive approach, to directly investigate nociceptors on molecular level at the neuro-cutaneous unit, which may have major implications for differential diagnostics and for the determination of potential therapeutic targets for analgesic and neuroprotective treatment approaches.

In conclusion, we optimized a method initially used for the isolation of synaptosomes from CNS to obtain isolated nerve endings from the mouse epidermis. The method was further validated by a translational approach on human skin punch biopsy. We applied this method to the well-studied Na_V_1.9 KO mouse model and identified pathways important for pain. This strategy can be further translated into both mouse pain models and human pain syndromes and help to understand the local proteomic and transcriptomic alterations in peripheral nerve branches associated with pain chronification and resolution.

## Methods

### Animals and ethical approval

The experiments on mice in this study adhered to the animal protection regulations specified by German federal law and the standards of the Association for Assessment and Accreditation of Laboratory Animal Care. These experiments received approval and oversight from the local veterinary authority. The mice were housed in the animal facility at the Institute of Clinical Neurobiology, University Hospital of Wuerzburg, under controlled conditions. C57BL/6 mice, Nav1.8cre::CAGeGFP, and Nav1.9 knockout mice were kept on a 12-hour day/night cycle, at temperatures between 20–22°C and humidity levels of 55–65%, with adequate food and water provided.

### Skin punch biopsy collection

We obtained 6-mm skin punch biopsies from the right lateral lower leg of n=4 healthy subjects (4 women) according to a standardized procedure [54].Biopsies were performed under sterile conditions and after local anesthesia (1 ml, 10 mg/ml Mecain, Puren Pharma GmbH, Munich, Germany). A commercial punch device was utilized (Kai Medical, Seki, Japan). Skin samples were collected in 8°C cold, sterile, DMEM/F-12 medium (Thermo Fisher Scientific, Waltham, MA, USA, 11320-074). Afterwards, the epidermis was manually isolated from each biopsy sample with a scalpel and directly transferred into ice-cold homogenization buffer (4 mM HEPES, 0.32M Sucrose, 1x Halt™ Protease Inhibitor Cocktail (Thermo Fisher Scientific, 78430), 40U/ml RNAsin (Promega) for further processing. Our study was approved by the Würzburg Medical School Ethics Committee (#49/23).

### Peripheral nerve isolation from mouse footpads and human epidermis

The epidermis from the footpads of P28 mice and human biospies were transferred into an ice-cold homogenization buffer (4 mM HEPES, 0.32M Sucrose, 1x Halt™ Protease Inhibitor Cocktail (Thermo Scientific, 78430), 40U/ml RNAsin (Promega, N2515). A glass tissue homogenizer was used with 12 gentle up-downs in 1:8 (w/v) of ice-cold homogenization buffer. The total homogenate (TH) was centrifuged at 1000 g for 10 min at 4 °C (Eppendorf 5417R). The supernatant (S1) is removed carefully from the pellet (P1) and centrifuged at 14600 x g for 15 min at 4 °C. The supernatant (S2) is removed completely, and the resulting fraction is referred to as the peripheral nerve fraction. The same procedure was applied to human epidermal tissues.Aliquots of all fractions were stored at −80 °C until further use.

### Immunohistochemistry

The epidermis from the hind paw of P28 mice was dissected in fixed in 4% PFA for 2h at room temperature. The post-fixed epidermis was embedded in 6% agarose in PBS. 20 µm of transverse sections were made using a vibratome (Leica VT1000S). For permeabilization and blocking, sections were first washed 3 times with PBS and blocked with 5% donkey serum, 0.3% Triton X-100, and 0.3% Tween 20 in PBS for 30 min at room temperature. For primary antibody labelling, the sections were incubated in antibodies for 48h at 4 °C in a blocking solution and immunolabeled with chicken anti-Nefh (1:5000, EMD Millipore, AB5539), mouse anti-Syn1 (1:1000, SYSY, 106 011), rabbit anti-Psd95 (1:1000, Abcam, ab18258), chicken anti-GFP (1:5000, Abcam, ab13970), and rabbit anti-PGP9.5 (1:10000, Proteintech, 14730-1-AP). The sections were washed 3 times with washing buffer (0.1% Triton X-100 and 0.1% Tween 20 in PBS) for 5 min each. The sections were stained with secondary antibodies donkey anti-chicken (1:10000, Jackson ImmunoResearch, 703-035-155), donkey anti-rabbit IgG-HRP (Jackson ImmunoResearch, 711-035-152), and donkey anti-mouse IgG-HRP (Jackson ImmunoResearch, 715-175-150) for 1h at room temperature and washed 3 times with washing buffer and 2 times with PBS. The sections were stained with 4′,6-diamidino-2-phenylindole (DAPI, Sigma-Aldrich, D9542) mounted on the objective slides, and embedded in FluorSave Reagent (Merck, 345789).

### Immunocytochemistry of peripheral nerve terminals

For peripheral nerve terminals antibody labelling, the protocol from Revelo et al. [41] was followed with a few modifications. The peripheral nerve terminals were immobilized on coverslips pre-coated with 5% BSA. For fixation, 4% PFA in PBS was used and was incubated at 4°C for 15 min on ice and 45 min at room temperature. Then, the peripheral nerve terminals were washed with PBS and quenched with 100 mM NH_4_Cl for 30 min. Peripheral nerve terminals were permeabilized and blocked using 0.1% Triton x-100 and 5% BSA in PBS. For primary antibody labelling, the peripheral nerve terminals were incubated in antibodies overnight at 4 °C in a blocking solution. Then the peripheral nerve terminals were washed 3 times with PBS and stained with secondary antibodies for 1h at room temperature and washed 3 times with PBS. The following primary and secondary antibodies were used: chicken anti-Nefh (1:5000, EMD Millipore, AB5539), mouse anti-Syn1 (1:1000, SYSY, 106 011), rabbit anti-Psd95 (1:1000, Abcam, ab18258), chicken anti-GFP (1:5000, Abcam, ab13970), rabbit anti-Rpl24 (1:1000, Thermo Scientific, PA5-30157), mouse anti-Rps6 (1:100, Thermo Scientific, MA5-15123), mouse anti-Syp1 (1:1000, SYSY, 101 011), mouse anti-Tomm20 (1:100, Santa Cruz Biotechnology, SC-17764), rabbit anti-PGP9.5 (1:10000, Proteintech, 14730-1-AP), donkey anti-chicken Alexa488 (Jackson ImmunoResearch, 703-545-155), donkey anti-mouse Cy3 (Jackson ImmunoResearch, 715-165-150), and donkey anti-rabbit Cy5 (Jackson ImmunoResearch, 711-175-152). Then, the coverslips were mounted on the objective slides, and embedded in FluorSave Reagent (Merck, 345789). The samples were imaged with an Olympus Fluoview 1000 confocal microscope.

### mCLING assay

For mCLING assay, the protocol from Ramirez et al. [55] was followed with a few modifications. The peripheral nerve terminals were incubated with 0.1µM mCLING-ATTO 647N-labeled (Synaptic Systems, 710 006AT1) for 10 min at room temperature. Then, a pulse of 70 mM KCl for 5 min was given to stimulate them. The peripheral nerve terminals were then fixed with 4% PFA and 0.2% glutaraldehyde in PBS for 15 min on ice and 45 min at room temperature. Then, the peripheral nerve terminals were washed with PBS and quenched with 100 mM Glycine in PBS for 5 min and 0.75 mM bromophenol blue for 5 min and washed 3 times with PBS. Peripheral nerve terminals were stained with primary antibodies as mentioned earlier for 1 h. Then the peripheral nerve terminals were washed 3 times with PBS and stained with the following secondary antibodies for 1h at room temperature and washed 3 times with PBS. Then, the coverslips were mounted on the objective slides, and embedded in FluorSave Reagent. The samples were imaged with an Olympus Fluoview 1000 confocal microscope.

### Protein extraction and western blotting

For peripheral nerve terminals, an equal volume of each fraction was diluted with 5× Laemmli buffer (250 mM Tris HCl pH 7.5, 5% SDS, 30% Glycerol, 5% β mercaptoethanol, 0.02% Bromophenol blue), heated at 99°C for 10 min and subjected to SDS-PAGE followed by transfer onto a nitrocellulose membrane (Transfer membranes GE Amersham™ Protran™, NCA2). The membrane was probed with following primary antibodies: Psd95 (Abcam, ab18258), Ck8 (Abcam, ab53280), Syp1 (SYSY, 101 011), PGP9.5 (Proteintech, 14730-1-AP), Histone H3A (Abcam, ab1791-100), GFP (Abcam, ab13970) for overnight at 4 °C and incubated with secondary antibodies like donkey anti-Rabbit IgG-HRP (Jackson ImmunoResearch, 711-035-152), donkey anti-mouse IgG-HRP (Jackson ImmunoResearch, 715-175-150), donkey anti-chicken (Jackson ImmunoResearch, 703-035-155) for 1h and developed using Thermo Scientific™ Pierce™ ECL kit (PI32106).

### Transmission electron microscopy

The isolated peripheral nerve endings were collected on the Isopore polycarbonate membrane filters (Merck, GTTP04700). The excess supernatant was vacuum-filtered. The filter was then incubated with preheated synaptosome regeneration buffer (H_2_O, 64 mM NaCl, 4 mM KCl, 0.8 mM CaCl_2_, 0.8 mM MgCl2 8 mM Tris-HCl (pH 7.4), 160 mM Sucrose) (37 °C) at room temperature for 20 min. The regeneration filter was removed again by vacuum filtration. The filter was immediately fixed in 2.5% glutaraldehyde in cacodylate buffer for 30 minutes on ice. The filter was washed with 50 mM cacodylate buffer three times, 10 minutes each on ice. The filter was then stored at 4 °C overnight [4]

For the whole epidermis, mice were deeply anesthetized and transcardially perfused with 0.1 M phosphate buffer (PB, pH 7.4) and 2.5% glutaraldehyde in cacodylate buffer. The epidermis was dissected and post-fixed for 90 min with 2.5% glutaraldehyde (50 mM cacodylate pH 7.2, 1 M KCl, 0.1 M MgCl_2_ at room temperature. The epidermis was washed with 50 mM cacodylate buffer three times, 10 minutes each.

The next steps were followed similarly for both the peripheral nerves on the filter and the mouse epidermis. The samples were fixed for 2 hours at 4°C with 2% OsO_4_ buffered with 50 mM cacodylate (pH 7.2), washed with H_2_O, and incubated overnight at 4°C with 0.5% uranyl acetate (in H_2_O). The samples were dehydrated, embedded in Epon812, and ultrathin longitudinal sectioned [39]. Electron micrographs were recorded with a JEOL JEM-2100 transmission electron microscope with a TVIPS F416 camera and a JEOL JEM-1400 Flash transmission electron microscope with a Matataki camera.

### Sample preparation for MS

Samples subjected to quantitative mass spectrometry analysis were first alkylated and reduced using 10 mM Tris(2-carboxyethyl)phosphine (TCEP), 40 mM 2-Chloracetamide (CAA) and 100 mM Tris-HCL pH 8.5 in 1% (w/v) sodium deoxycholate (SDC) at 45 °C for 5 min. Digestion was then done with a 1:1000 protein to LysC and Trypsin (1:1) ratio overnight with 1400 rpm agitation on an Eppendorf Thermomixer C at 37 °C. For desalting peptides were first diluted tenfold using 1% trifluoroacetic acid (TFA) in isopropanol and loaded on SDB-RPS (Empore) StageTips [40]. The flowthrough was discarded and StageTips were washed once with 200 mL of 1% TFA in isopropanol and then twice with 0.2% TFA/2% acetonitrile (ACN). Peptides were finally eluted using 60 mL of 80% ACN/1.25% NH_4_OH and dried using a SpeedVaccentrifuge (ConcentratorPlus; Eppendorf) for 1 hour at 30 °C. Peptides were then resuspended in 0.2% TFA/2% ACN. 200 ng of peptides were used for LC-MS/MS analysis.

### Data-independent acquisition

For the data-independent acquisition, we used the following MS setup: 50 cm reversed-phase column (75 µm inner diameter, packed in-house with Reprosil-Pur C18-AQ 1.9 µm resin) that was kept at 50°C using a homemade column oven. The mass spectrometer (Orbitrap Exploris 480, Thermo Fisher Scientific) was connected online with an EASY-nLC 1200 system (Thermo Fisher Scientific). For peptide separation, we used a binary buffer system consisting of 0.1% formic acid (FA) (buffer A) and 80% ACN, 0.1% FA (buffer B). We used a constant flow rate of 300 nL/min and a 75 min gradient. The gradient starts with 2% buffer B and gradually increases to 35% within 60 min, 60% within 70 min and 90% within 71 min, and stays constant until 75 min. We used the following MS settings: data-independent acquisition (DIA) mode with a full scan range of 300–1650 m/z at resolution of 120,000, a maximum injection time of 20 ms, automatic gain control of 3e6, with a stepped higher-energy collision dissociation (HCD) (set to 25, 27.5, and 30). After each full scan, 44 DIA scans were followed with a 30,000 resolution, an AGC of 1e6, and a maximum injection time of 54 ms [46].

### DIA data processing and data analysis

DIA-NN [9] 1.8.1 was used as a search engine to process DIA raw files. Mus musculus and a human reference proteome with canonical and isoform sequences were used as the FASTA file. For the search, default settings were used with the following exceptions: For protein modification, we included carbamidomethylation, oxidation of methionine, and N-terminal acetylation. FASTA digest for library-free search and the Deep learning-based spectra, RTs, and IMs predictions including the heuristic protein inference was turned on. Data analysis and visualization were done using Python v3.5.5 with the following libraries: seaborn 0.11.2, pandas 1.4.2, scikit-learn 1.5, numpy 1.21.5, matplotlib 3.5.13, statsmodels 0.13.5 and gseapy 1.0.4. Protein intensities were the first log2 transformed to approximate a normal distribution. Data was then filtered for valid values in at least one experimental group.

Imputation was done using a sampling method from a shifted Gaussian distribution with a shift of 3 and width of 0.3 standard deviations. Hierarchical clustering was performed using the euclidean distance. For comparison of S2 fraction from wild type and Na_V_1.9 KO, quadruplicates were analyzed and one outlier was removed. For determining statistical significance we used an unpaired two-tailed Student’s t-test. To address false positives we adjusted the p-values using the Benjamini-Hochberg method. Enrichment analysis was done using the GSEAPY tool [17].

### Image acquisition and data analysis

Images were acquired using an Olympus Fluoview 1000 confocal system equipped with the following objectives: 10× (NA: 0.25), 20× (NA: 0.75), 40× (oil differential interference contrast, NA: 1.30), and 60× (oil differential interference contrast, NA: 1.35). Fluorescence excitation was achieved with 405, 473, 559, and 633 nm lasers. Visualization was done using the Olympus FV10-ASW (RRID: SCR_014215) imaging software. Structured illumination microscopy (SIM) was conducted with a commercial Zeiss Elyra S.1 microscope, featuring a Plan-Apochromat 63× NA 1.40 immersion-oil-based objective and four excitation lasers: a 405 nm diode (50 mW), a 488 nm OPSL (100 mW), a 561 nm OPSL (100 mW), and a 642 nm diode laser (150 mW). Imaris software (Oxford Instruments) was used for 3D reconstruction, and the resulting images were processed by maximum intensity projection and adjusted for brightness and contrast using Image J software.

### Statistics and reproducibility

All statistical analyses were performed using GraphPad Prism version 9 for Windows (GraphPad Software, San Diego, California USA) and statistical significance was considered at P<0.05. Quantitative data are presented as mean ± s.d. unless otherwise indicated. No statistical method was used to pre-determine the sample size. No data were excluded from the analyses. Two groups were compared using unpaired two-tailed Student’s t-tests or two-tailed one-sample t-tests. For multiple independent groups, one-way analysis of variance (ANOVA) with post hoc multiple comparisons test was used. Details of replicate numbers, quantification and statistics for each experiment are specified in the Figure legends.

## Acknowledgments

We thank Prof. Claudia Sommer and Prof. Esther Asan for their assistance in the annotation of specific cell types in whole epidermal electron micrographs. We also thank Prof. Markus Sauer for the SIM (Zeiss Elyra S.1). The study was supported by the German Research Foundation (DFG) Clinical Research Group ResolvePAIN KFO5001 (MB4910/3-1 to M.B. and SE697/9-1 to M.S.). The JEOL JEM-2100 transmission electron microscope is funded by the DFG grant 218894163, and the JEM-1400 Flash transmission electron microscope is funded by DFG grant 426173797 (INST 93/1003-1 FUGG).

## Conflict of interest statement

The authors have no conflicts of interest to declare.

## Supplementary figure legends

**Supplementary Figure 1.** Transmission electron microscopy of whole epidermis and isolated peripheral nerve terminal

(A) Transverse section of whole mouse epidermis showing unmyelinated peripheral nerve terminal (red arrowheads), and keratinocyte (K). Scale bar: 0.2 nm. (B) Electron micrograph of isolated peripheral terminals following fractionation immobilised on polycarbonate filter. Scale bars: 200 nm.

**Supplementary Figure 2.** Immunohistochemistry of Na_V_1.8cre::CAGeGFP mouse epidermis

Confocal image of mouse epidermis from control and Na_V_1.8cre::CAGeGFP mice stained with GFP, PGP9.5, and DAPI show an overlap between GFP labelled Na_V_1.8 fibers. Scale bar: 50 µm.

**Supplementary Figure 3**

PCA and cluster mapping of the S2 fraction from wild type and Na_V_1.9 KO mice before (A and B) and after the outlier removal (C and D).

**Supplementary Figure 4.** Gene ontology for differentially expressed proteins

(A and B) Dot plot showing GO terms related to molecular functions for upregulated (A) and downregulated (B) proteins, respectively, in peripheral nerve terminals of wild type vs Na_V_1.9 KO mice. (C and D) GO terms related to cellular components for upregulated (C) and downregulated (D) proteins respectively in peripheral nerve terminals of wild type vs Nav1.9 KO mice.

**Supplementary Fig. 5**

STRING analysis of upregulated proteins in peripheral nerve terminals of wild type relative to Na_V_1.9 KO mice. Only interactions with the highest confidence score (0.9) are shown.

**Supplementary Fig. 6**

STRING analysis of downregulated proteins in peripheral nerve terminals of wild type relative to Na_V_1.9 KO mice. Only interactions with the highest confidence score (0.9) are shown.

## Notes

### Competing Interest Statement

The authors have declared no competing interest.

