## Supplementary figures and images for "Proteomic analysis of isolated nerve terminals from Na_V_1.9 knockout mice reveals pathways relevant for neuropathic pain signalling"

### Supplementary Fig. 1

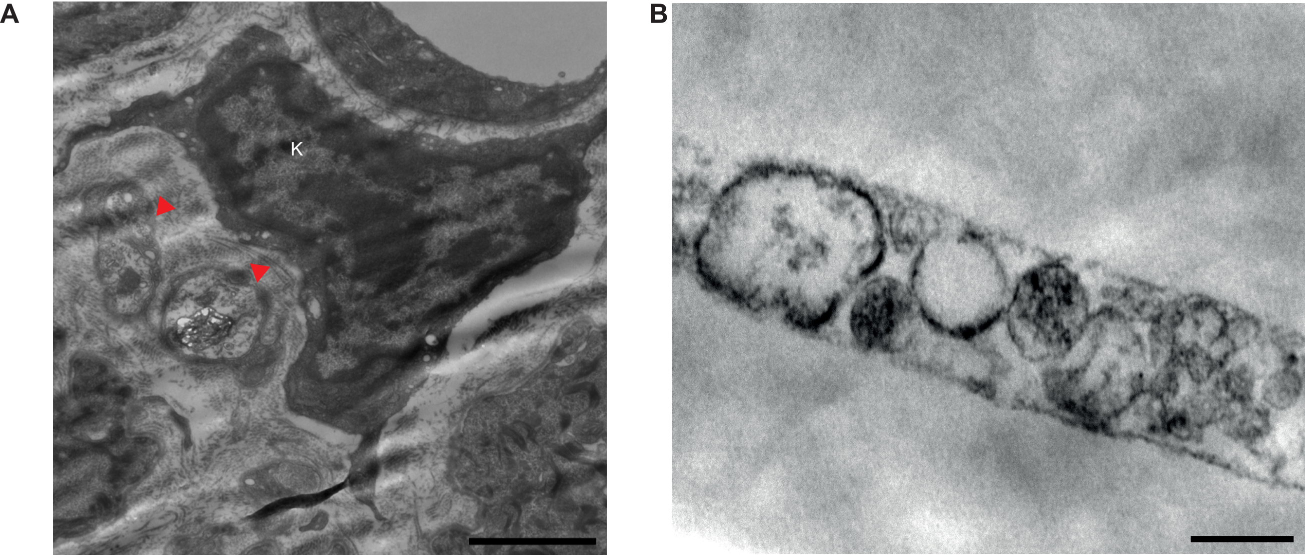

### Supplementary Fig. 2

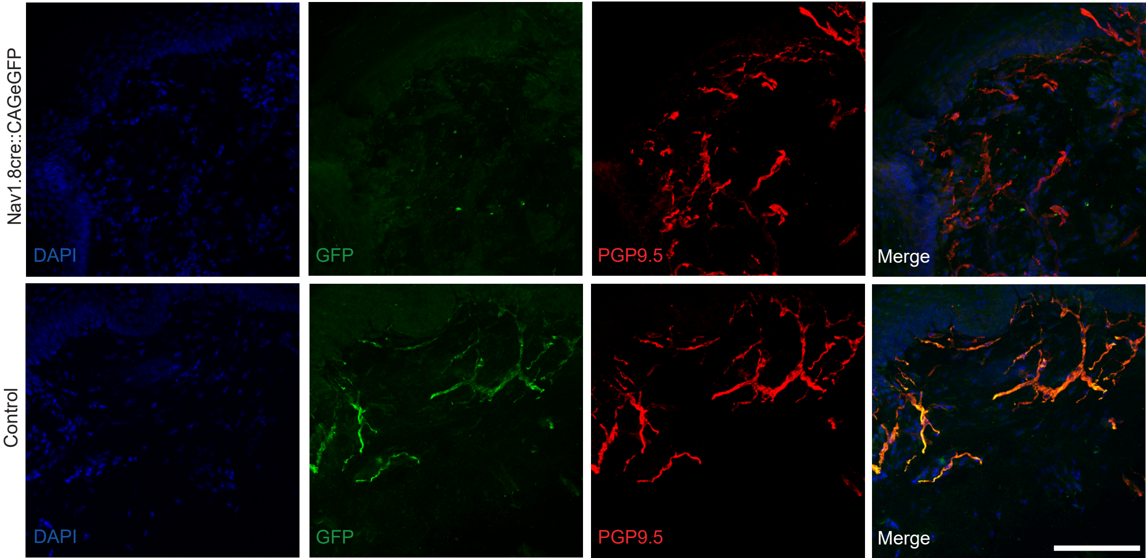

### Supplementary Fig. 3

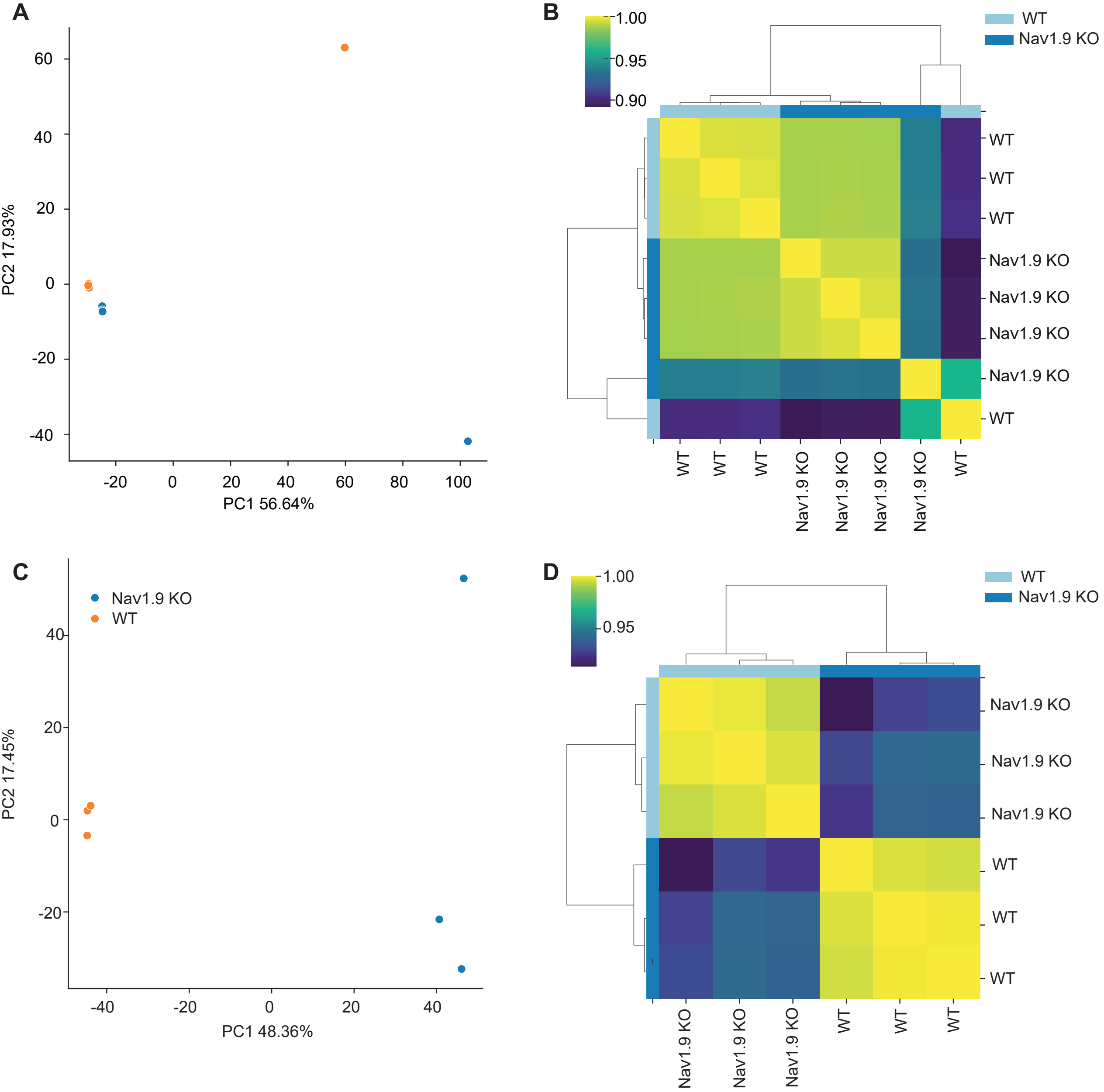

### Supplementary Fig. 4

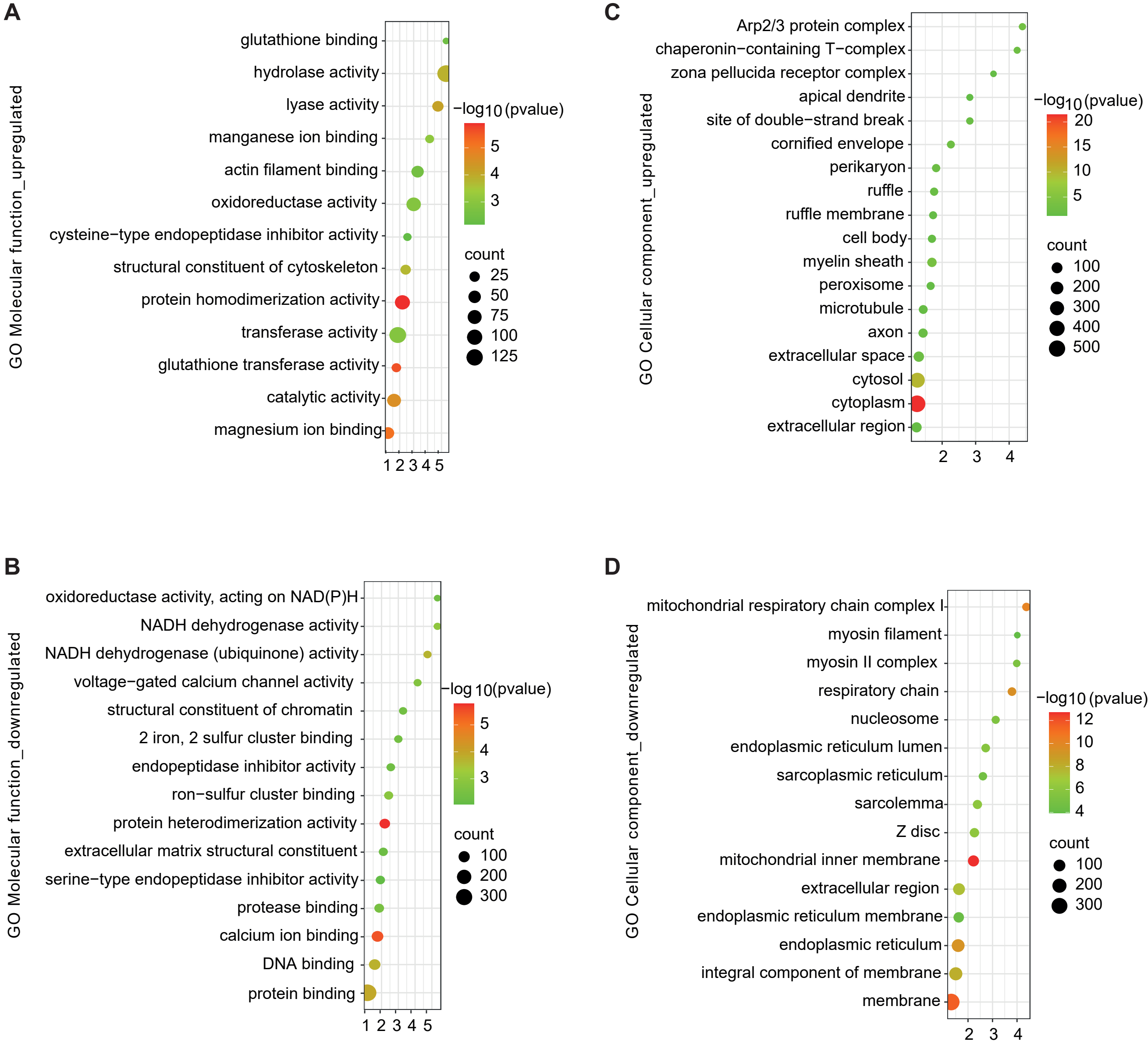

### Supplementary Fig. 5

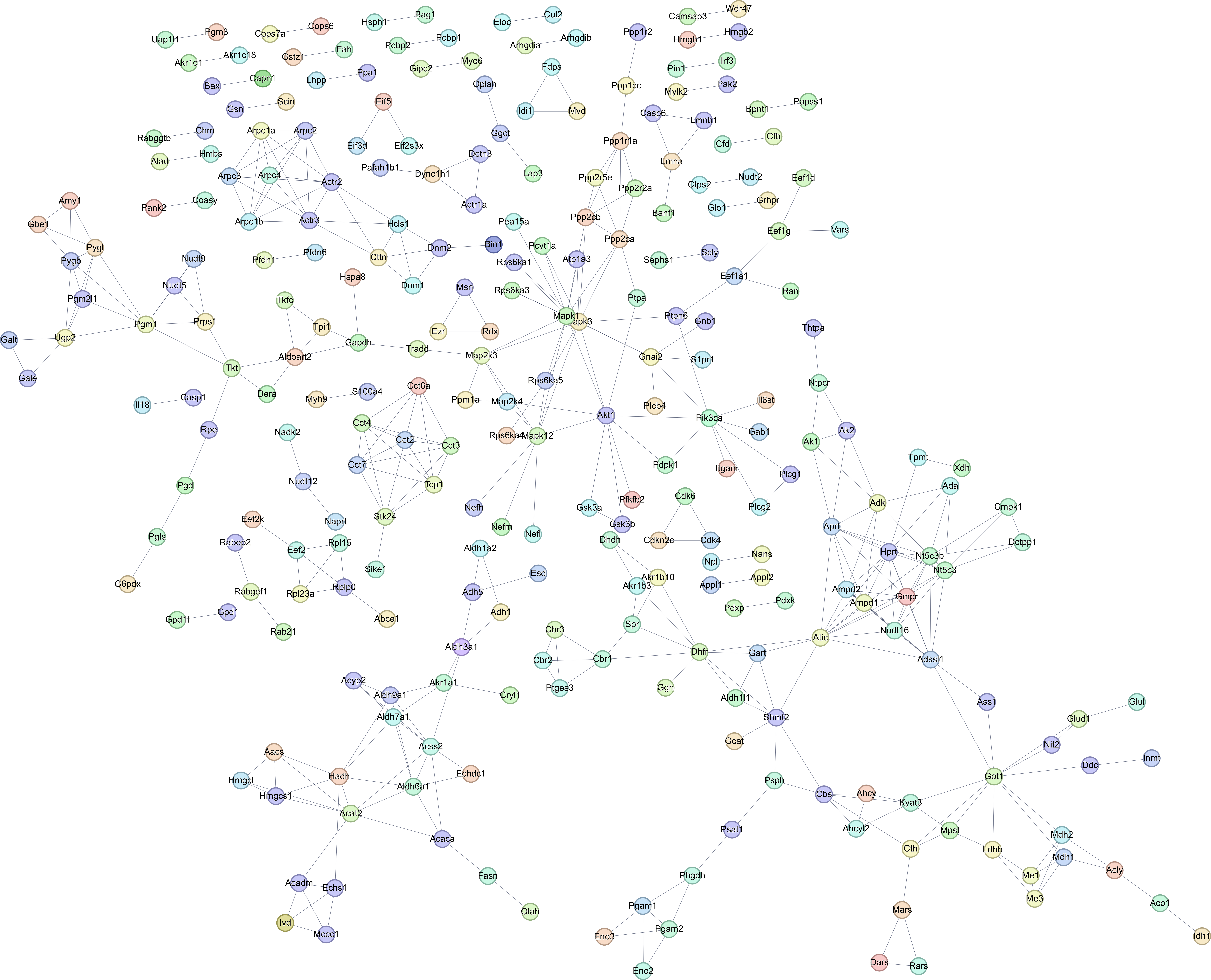

### Supplementary Fig. 6

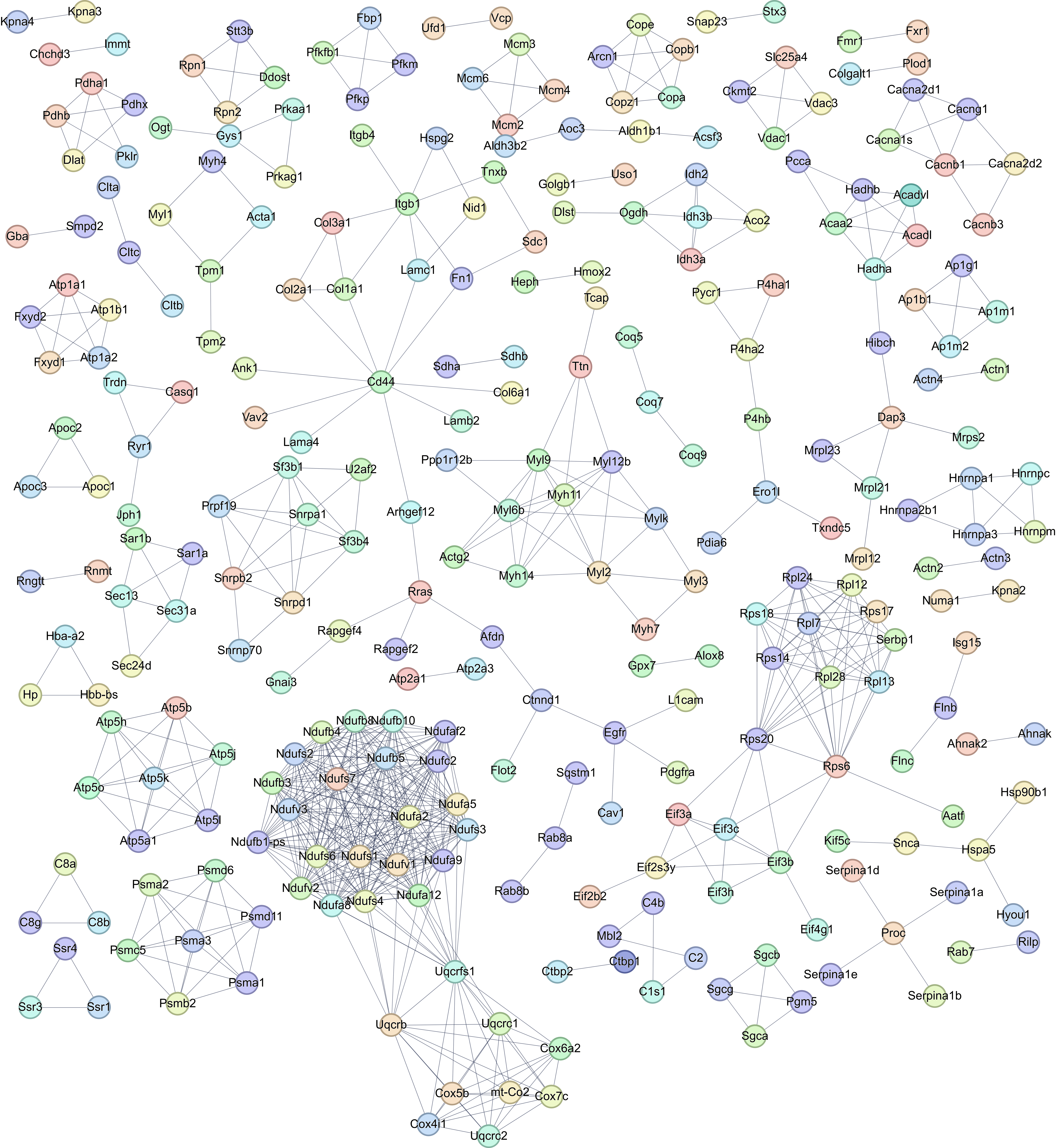
